# Genetically Engineered Cell Membrane-Coated Nanodrug for Targeted Treatment of Thyroid Cancer

**DOI:** 10.1101/2025.10.15.682623

**Authors:** Shaojie Xu, Xingyin Li, Youyun Peng, Xue Yang, Yuhang Su, Hanning Li, Sining Wang, Xingrui Li, Yaying Du

## Abstract

Thyroid cancer is the most common endocrine malignancy, particularly in patients with radioactive iodine-refractory differentiated thyroid cancer (RAIR-DTC), who have limited treatment options and poor clinical prognosis. In this study, doxorubicin (DOX) and sorafenib (SOR) were loaded into the photothermal conversion agent mesoporous polydopamine (mPDA), forming mPDS. Tumor cell membrane coating engineering resulted in the formation of mPDS@CAR-M, significantly improving tumor targeting. After internalization by tumor cells, the loaded drugs are rapidly released under near-infrared (NIR) laser irradiation. mPDS@CAR-M effectively inhibits tumor cell proliferation by enhancing oxidative stress, suppressing the PI3K-AKT-mTOR signaling pathway, and inducing cytotoxic autophagy. By activating harmful autophagy, mPDS@CAR-M further inhibits the epithelial-mesenchymal transition (EMT) process, reducing tumor cell migration capacity. In vivo experiments showed that mPDS@CAR-M significantly reduced tumor volume, with its therapeutic efficacy closely related to the expression level of the targeted surface antigen. Therefore, mPDS@CAR-M demonstrates significant potential in the treatment of RAIR-DTC, providing a novel direction for further clinical exploration.

## Introduction

Thyroid cancer is the most common endocrine malignancy, with a steadily rising global incidence[1]. In patients with locally advanced or metastatic differentiated thyroid cancer (DTC), approximately two-thirds eventually progress to radioactive iodine–refractory DTC (RAIR-DTC), for which therapeutic options remain limited and clinical outcomes are poor, with a 10-year survival rate of only 15%[2]. Although doxorubicin (DOX) and sorafenib (SOR) are FDA-approved chemotherapeutic and targeted agents for this disease respectively[3,4], their clinical utility has declined due to resistance and toxicity. Notably, the DOX–SOR combination has shown efficacy in hepatocellular carcinoma[5,6], but its role in thyroid cancer remains unverified.

Cell membrane–coated nanoparticles (CNPs) are fabricated by cloaking nanoparticle cores with cell membranes, thereby endowing them with the surface markers of the source cells. By modulating membrane protein expression, genetically engineered CNPs can reshape their biointerface and enhance functional performance. The use of CAR–T cell membrane–coated nanoparticles for tumor therapy has been validated in hepatocellular carcinoma models[7]. However, clinical-grade CAR–T cells typically exhibit low transfection efficiency and rely on terminally differentiated effector T cells[8]. In contrast, tumor cells are easily transfected and expanded, and possess intrinsic homotypic adhesion properties, making them an ideal cellular source for constructing CAR-engineered membranes.

Tumor-specific antigens (TSAs), expressed exclusively on the surface of tumor cells[9], represent ideal therapeutic targets. However, when tumor cells simultaneously present TSAs and CARs on their surfaces, the resulting membrane-coated nanoparticles may undergo self-aggregation and deposition, thereby compromising their targeting capacity and stability. The thyroid-stimulating hormone receptor (TSHR) is a compelling therapeutic target owing to its consistent expression in the majority of clinical RAIR-DTC and lymph node metastatic lesions[10], while its absence in most thyroid cancer cell lines renders it an ideal target antigen for constructing CAR-M.

In this study, we developed a genetically engineered, membrane-coated nanodrug (mPDS@CAR-M), in which the photothermal agent mesoporous polydopamine (mPDA) was employed to co-load DOX and SOR, followed by cancer cell membrane coating to achieve targeted delivery. Upon near-infrared (NIR) irradiation, the system enabled spatiotemporally controlled drug release through a photothermal effect, thereby achieving a synergistic combination of photothermal therapy, chemotherapy, and targeted therapy for effective tumor ablation. Experimental results demonstrated that mPDS@CAR-M enhanced therapeutic efficacy through autophagy modulation, offering a potential treatment strategy for patients with RAIR-DTC (Figure 1).

**Figure 1.**
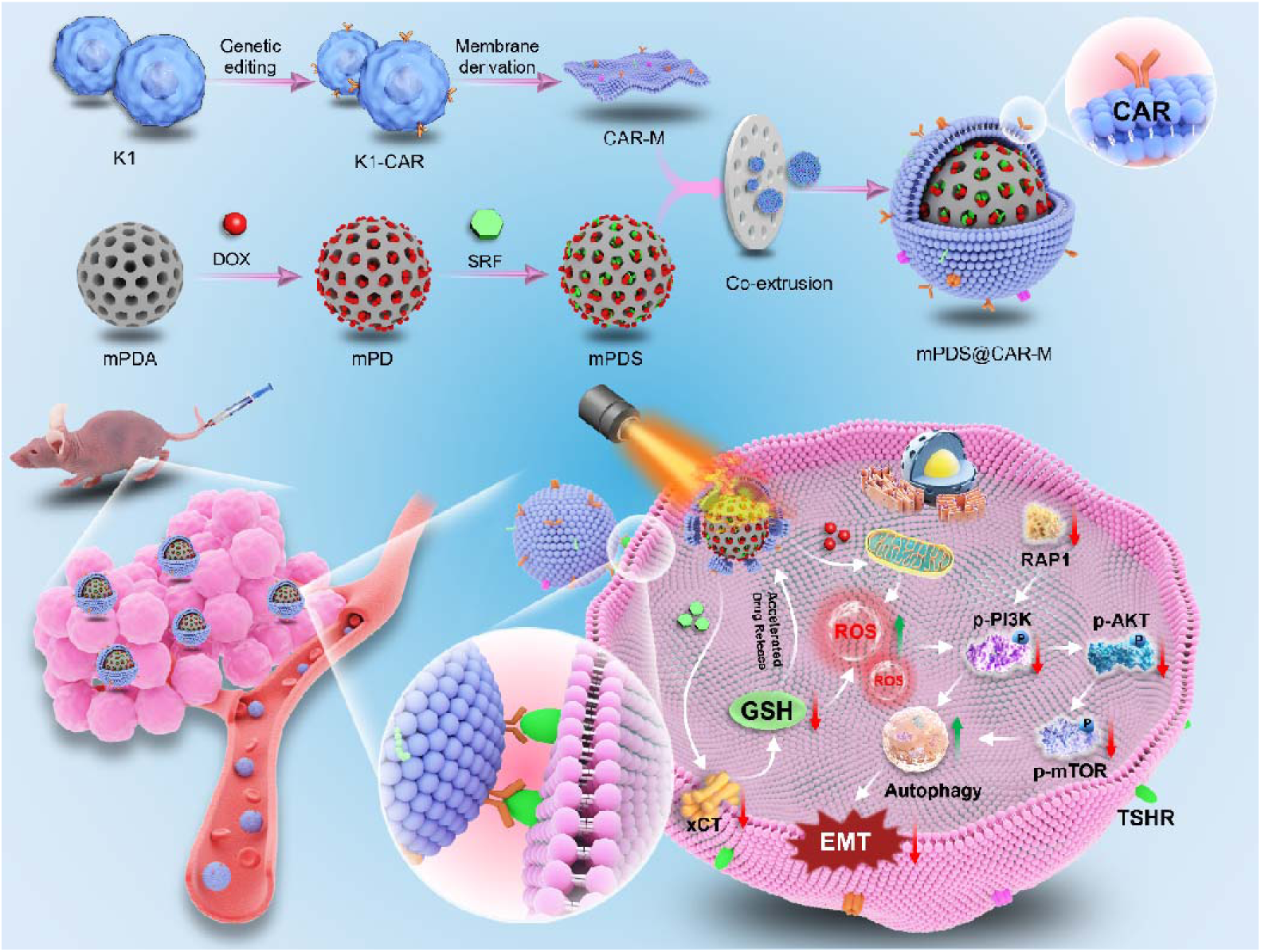
The synthesis and mechanism diagram of mPDS@CAR-M. mPDS@CAR-M targets thyroid cancer by interacting with the TSHR on the surface of thyroid cancer cells and promotes cellular damage-induced autophagy and inhibits EMT through the suppression of the PI3K-Akt-mTOR signaling pathway.

## Materials and Methods

### Establishment of stable cell lines

K1,IHH4,TPC-1,BCPAP,FTC-133 and Nthy-ori 3-1K1 cells were purchased from icell (Shanghai, China) and cultured in RPMI-1640 medium (Thermo Fisher, Waltham, MA, USA) supplemented with 10% FBS (Thermo Fisher, Waltham, MA, USA). K1 and IHH4 cells were transduced with lentiviruses carrying TSHR or TSHR-specific CAR sequences in serum-free medium. After 12 h, medium was replaced, and antibiotic selection was applied for 7–10 days to establish TSHR- and CAR-expressing stable lines. To generate cell lines with varying levels of TSHR expression, different viral loads were used during transduction. The cells were then sorted or selected based on their TSHR expression levels, as determined by flow cytometry, to obtain populations with low, medium, and high expression of TSHR.

### Flow Cytometry Analysis

For flow cytometric analysis, cells were harvested and washed with cold PBS. The cells were then incubated with the appropriate fluorescent dye, such as phycoerythrin (PE) for TSHR staining or DCFDA-DA for ROS detection, for an appropriate amount of time at 4°C in the dark. After washing with PBS, the cells were resuspended in PBS and analyzed using a flow cytometer. Fluorescence signals were collected, and data were analyzed using FlowJo to quantify the fluorescence intensity. A minimum of 10,000 events were recorded for each sample, and appropriate controls were used to set compensation and gating strategies.

### Synthesis and characterization

mPDA was purchased from Xi’an Ruixi Biologicals, while doxorubicin (DOX) and sorafenib (SOR) were from MedChemExpress. Nanoparticles were prepared by mixing 10 mg mPDA with 5 mg DOX, 5 mg SOR, or both, followed by overnight stirring, centrifugation, washing, and redispersion (yielding mPD, mPS, and mPDS). Tumor cell membranes were isolated from CAR-K1 and CAR-IHH4 cells using the Minute™ Plasma Membrane Kit. Extracted tumor cell membranes were sequentially extruded through 400nm and 200nm polycarbonate membranes for approximately 20 cycles to obtain CAR-M. mPDS@M and mPDS@CAR-M were prepared by co-extruding the nanoparticles with tumor cell membranes using the same procedure. mPDS@CAR-M, CAR-M, and mPDS solutions were sonicated for 5 minutes, separately applied to copper grids, and negatively stained with uranyl acetate for observation using a low-voltage transmission electron microscope (TEM) system (Hitachi, Japan). mPDS was dropped onto a silicon wafer, dried under an infrared lamp, and then attached to a sample holder with conductive adhesive for scanning electron microscopy (SEM) imaging (Hitachi, Japan). For cell samples, fixation with osmium tetroxide was followed by phosphate-buffered saline washing, dehydration, embedding, sectioning, staining, and air drying, after which the cellular uptake of nanoparticles could be observed under a TEM.

The particle size distribution and ζ potential of mPDA, mPDS, CAR-M, and mPDS@CAR-M solutions were analyzed by dynamic light scattering (DLS, Malvern-Zetasizer, UK). The particle size distributions of mPDS@CAR-M, mPDS, and mPDA were monitored over 48 hours, with absorbance at 540 nm measured at 1, 2, 4, 12, 24, and 48 hours after mixing with an equal volume of fetal bovine serum (FBS) and incubating at 37°C to assess the serum stability of the materials.

Additionally, hemolysis was evaluated by incubating mouse red blood cell suspensions with PBS (negative control), deionized water (positive control), or serially diluted mPDS@CAR-M solutions for 3 hours, followed by centrifugation and measurement of supernatant absorbance at 540 nm.

Co-extruded cell membrane vesicles and nanoparticles were analyzed by sodium dodecyl sulfate–polyacrylamide gel electrophoresis (SDS-PAGE). Briefly, protein samples were prepared by mixing with loading buffer, then heated at 100°C for 10min. The samples were loaded onto a polyacrylamide gel and subjected to electrophoresis. The protein bands were then stained with Coomassie Blue.

Powders of mPDS were mixed with KBr and pressed into transparent pellets for recording FTIR spectra in the range of 500–4000 cm-1 using an FTIR spectrometer (Thermo Fisher, USA). Aqueous solutions of the mPD,mPS,mPDS were placed in quartz cuvettes, and their absorbance at different wavelengths was measured with a UV–Vis spectrophotometer (Perkin-Elmer, USA). Standard curves for DOX and SOR were established by preparing solutions with defined concentration gradients. DOX absorbance was measured with a UV–Vis spectrophotometer, while the peak areas of SOR solutions were determined by high-performance liquid chromatography (HPLC). During mPDS synthesis, the supernatant and wash solutions were collected after three washes, and drugs content were quantified using the corresponding calibration curves. The drug loading efficiency (%) and the drug loading content (mg/mg) were calculated as follow:

Encapsulation Efficiency (%) = (Mused - MSupernatant - MWash) / MUsed × 100% Loading Efficiency (%) = (MUsed - MSupernatant - MWash) / (MAgent + MmPDA) × 100% where MUsed is the total amount (mg) of drug initially added, MSupernatant and MWash are the drug amounts (mg) in the supernatant and wash solution, MAgent is the weight(mg) of DOX or SOR encapsulated inside the nanoparticles and MmPDA is the total amount (mg) of mPDA used in nanoparticle synthesis.

The release of drugs was monitored at predetermined time intervals, both with and without near-infrared (NIR) irradiation, using the same detection methods.

Cumulative release percentage (%) was defined using the following formulas:

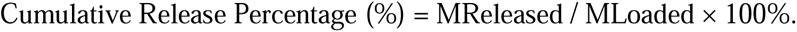

where MReleased represents the total weight (mg) of the released drug and MLoaded is the total weight (mg) of the initial loaded drug.

For evaluating the photothermal performance of the material, both in vitro and in vivo methods were employed. In vitro, mPDS@CAR-M was irradiated with an 808 nm NIR laser (2 W/cm²) in 2 min on/2 min off cycles (4 min per cycle) for three cycles, with temperature monitored by an infrared thermal imager (Testo 865; Testo, Schwarzwald, Germany). In vivo, mPDS@CAR-M was administered to nude mice via tail vein injection; on alternate day, the tumor site was irradiated with laser for 2 min, with local temperature monitored and recorded.

### Cell functional assays

K1-TSHR cells were treated with 10 μg/mL of PBS, mPS, mPD, mPDS, mPDS@M, or mPDS@CAR-M for 12, 24, and 48 h, or with 5, 10, or 20 μg/mL for 48 h, and cell viability was assessed by the CCK-8 assay. The live/dead, migration, and colony formation assays used the same treatment groups as the CCK-8 assay. Live/dead staining was performed after treatment with 10 μg/mL for 48 h using a Calcein-AM/propidium iodide kit. For migration and colony formation, cells were pretreated at 10 μg/mL for 48 h, then processed as follows: for migration, pretreated cells were resuspended in 100 μL serum-free medium and seeded into the upper chambers of Transwell inserts (lower chambers, 10% FBS) and, after 8–12 h, migrated cells were fixed and stained; for colony formation, pretreated cells were reseeded at 800 cells per well in six-well plates, the medium was changed every 3 days, and colonies were fixed, stained, and counted. The same procedures were applied to TSHR-IHH4 cells.

K1-TSHR cells were treated with mPDS@CAR-M (0, 2, 5, 10, and 20 μg/mL) for 48 h. ROS levels were measured using DCFH-DA staining followed by flow cytometry. For glutathione detection, cell lysates were prepared using Protein Removal Reagent M, followed by freeze-thaw and centrifugation. The absorbance at 412 nm was measured using a microplate reader. Absorbance values for the standards were measured in parallel, and a standard curve was constructed. The total glutathione and GSSG content in the samples were calculated based on the standard curve.

### Immunofluorescence staining

Equal numbers of K1 and K1-TSHR cells were seeded in 24-well plates and treated with DiD-labeled CAR-M. Immunofluorescence (IF) was used to assess the targeting specificity of CAR-M to K1 and K1-TSHR cells. K1-TSHR cells were seeded in 24-well plates and treated with equal amounts of DiD-labeled M and CAR-M. IF was used to evaluate the targeting properties of M and CAR-M to K1-TSHR cells.

mPDS was labeled with FITC (green), the membrane was labeled with DiD (red), and the nuclei were stained with DAPI (blue). After co-extrusion, dual-fluorescently labeled mPDS@M and mPDS@CAR-M were obtained. K1-TSHR cells were seeded in confocal dishes (3 × 10 cells/well) and cultured for 12 h. Subsequently, 10 μL of mPDS@M or mPDS@CAR-M solution at equal concentrations was added to the cells in FBS-free medium and incubated for 3 h. Images were acquired using confocal laser scanning microscopy (CLSM) with the following excitation/emission parameters: DiD (Ex/Em: 644/663 nm) and FITC (Ex/Em: 492/518 nm).

FITC-labeled mPDS, PDS@M, and mPDS@CAR-M were intravenously injected via the tail vein into tumor-bearing nude mice. At 24 h post-injection, the biodistribution of nanoparticles was assessed using an in vivo imaging system. After euthanasia, the heart, liver, spleen, lungs, kidneys, and tumor tissues were harvested, rinsed with physiological saline, and imaged for fluorescence using an animal imaging system.

### Animal Models

BALB/c Nude mice (male, 3–4 weeks) were obtained from Jiangsu GemPharmatech (Jiangsu, China) and maintained under SPF conditions. All procedures were approved by the Animal Ethics Committee of Tongji Hospital, Tongji Medical College (Huazhong University of Science and Technology, Wuhan, China).

To evaluate the therapeutic efficacy of membrane-coated nanoparticles in a K1-TSHR subcutaneous tumor model, male BALB/c nude mice were inoculated subcutaneously with K1-TSHR tumor cells (3–4 × 10 cells in 100 μL). When the tumors reached approximately 6mm in diameter, the mice were randomly divided into six groups: control, mPS, mPD, mPDS, mPDS@M, and mPDS@CAR-M. Different drugs at the same concentration were administered via tail vein injection. At 24 h post-injection, the tumor sites were irradiated with an 808 nm NIR laser (2 W/cm²) for 2 min, repeated three times. Tumor volume and body weight were monitored daily. On day 18, all mice were euthanized, and blood was collected via the orbital vein. Tumors, hearts, livers, spleens, lungs, and kidneys were collected. The levels of ALT, AST, CK-MB, and LDH1 in the blood of nude mice from different groups were measured. Tumors were photographed and weighed, and both tumors and organs were subjected to paraffin embedding, sectioning, and histological analysis. Organs were stained with hematoxylin and eosin (H&E). Additionally, tumor sections were stained with H&E and immunohistochemistry (IHC) for Ki67, CD31, LC3, and p-AKT.

Further, male BALB/c nude mice were randomly divided into four groups for transplantation of different tumor cells (K1-TSHR+, K1-TSHR++, K1-TSHR+++, and wild-type K1). All four groups received the same mPDS@CAR-M treatment and NIR irradiation as described above, with drug injections carried out at 3-day intervals between each treatment and tumor size was monitored daily. On day 20, the experiment was terminated, mice were euthanized, and tumors were collected for photographing and weighing.

### RNA Sequencing

K1-TSHR cells treated with 10μg/mL mPDS@CAR-M for 48 h were collected for transcriptome sequencing on the Illumina platform: Total RNA was extracted, cDNA libraries were constructed and purified, and RNA-seq was performed for in-depth analysis. Three biological replicates per group were included.

### Reverse transcriptase quantitative polymerase chain reaction (RT-qPCR)

K1-TSHR cells treated with 10 μg/mL mPDS@CAR-M for 48 hours, as well as those treated with PBS, were collected. Total RNA was extracted using the RNeasy Mini Kit, and cDNA was synthesized using the iScript cDNA Synthesis Kit. RT-qPCR was performed using SYBR Premix Ex Taq and the MxPro Mx3005P real-time PCR detection system (Agilent Technologies, Santa Clara, CA). Primer sequences are listed in Supplementary Table 1. GAPDH was used as an internal control.

### Western blot

Protein expression levels were assessed by Western blotting and quantified using ImageJ. Briefly, K1-TSHR cells from each group were collected after 48 h of treatment and lysed on ice for 20 min in 1× RIPA buffer supplemented with 1% PMSF. Protein concentrations were determined using a BCA assay kit (Boster, Hubei, China). Lysates were denatured at 100 °C for 10 min, and 20 μg of protein per lane was separated on SDS–PAGE and transferred to PVDF membranes. Membranes were blocked with a rapid blocking buffer at room temperature for 15–30 min, incubated with appropriately diluted primary antibodies overnight at 4 °C on a shaker, washed with TBST, and then incubated with species-specific secondary antibodies for 30–60 min. Immunoreactive bands were visualized using an enhanced chemiluminescence (ECL) detection kit.The primary antibodies included TSHR, APT1A1, xCT, RAP1A, mTOR, p-mTOR, PI3K, p-PI3K, AKT, p-AKT, SQSTM, LC3- / , GAPDH and β-tubulin (Abclonal, hubei,china). Mouse anti-human β-actin and goat anti-mouse/anti-rabbit secondary antibodies were purchased from Proteintech.

### Statistical Analyses

The data were analyzed using t tests, one-way ANOVA or Kruskal–Wallis test in Prism 9 (GraphPad, La Jolla, CA, USA). p < 0.05 was considered to indicate significance.

## Result and Discussion

The expression of TSHR in different thyroid cancer cell lines remains controversial[10]. We first confirmed that TPC-1, K1, IHH4, BCPAP, and Nthy-ori 3.1 cells completely lacked TSHR expression, whereas FTC133 cells expressed TSHR (Supplementary Figure 1A). To further investigate, we selected the K1 cell line derived from a primary tumor and the IHH4 cell line derived from lymph node metastatic lesions, and established two stable TSHR-expressing cell models, K1-TSHR and IHH4-TSHR, via lentiviral transduction. TSHR was successfully expressed and localized to the plasma membrane, as confirmed by flow cytometry (Figure 2A), qPCR (Figure 2C), and Western blotting (Figure 2D). The construction of CAR-K1 cells followed a similar procedure to that used for our previously established CAR-T cells[11], and CAR expression on the cell surface was confirmed by flow cytometry (Figure 2B).

**Figure 2.**
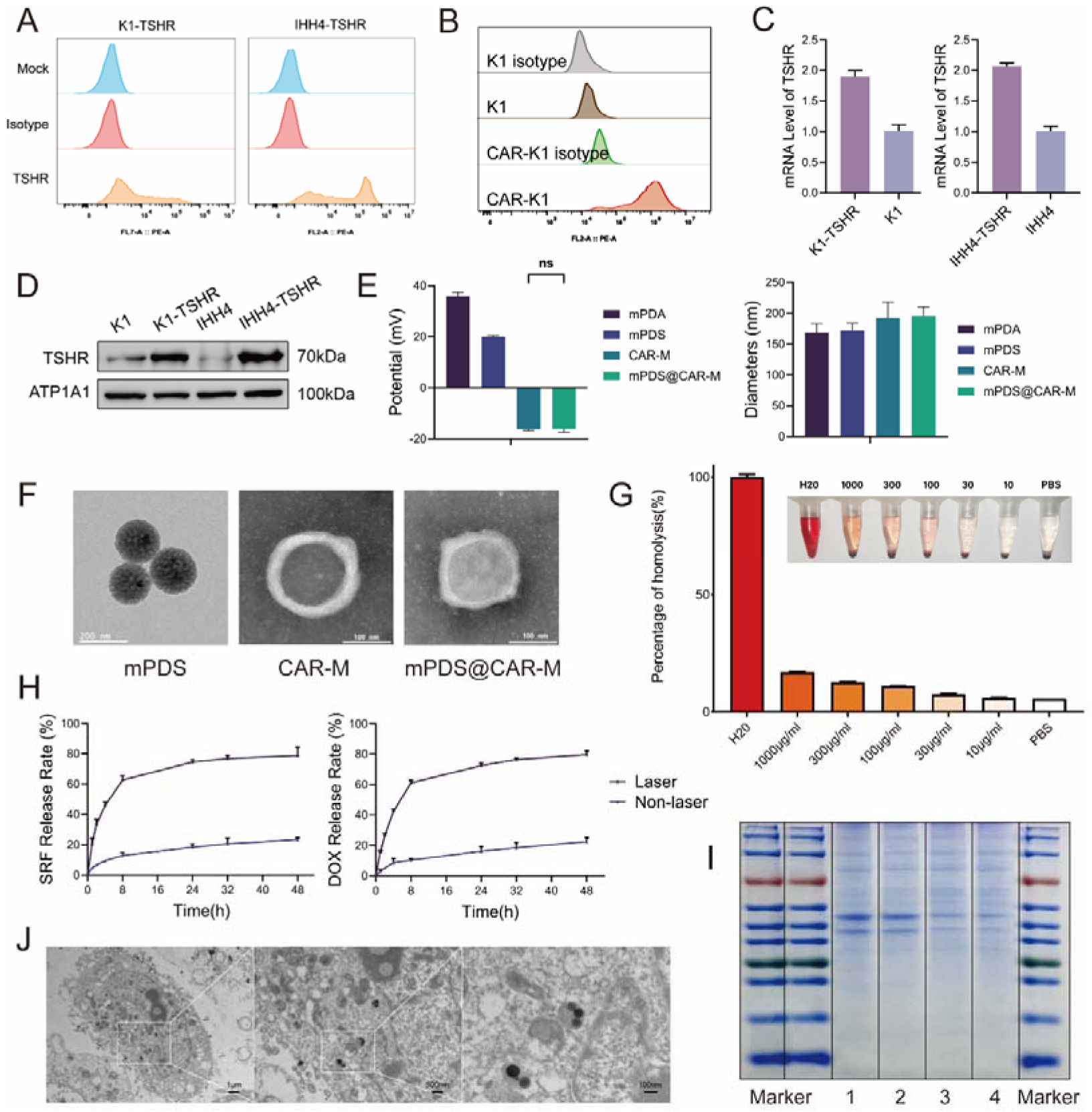
Characterizations of mPDS@CAR-M. (A) Flow cytometry analysis of TSHR expression in K1-TSHR and IHH4-TSHR cells. (B) Flow cytometry validation of CAR expression on the surface of CAR-K1 cells. (C) qPCR analysis confirming TSHR expression in K1-TSHR and IHH4-TSHR cells. (D) Western blot analysis verifying TSHR expression on the cell membranes of K1-TSHR and IHH4-TSHR cells. (E) Zeta potential and particle size measurements of mPDA, mPDS, CAR-M, and mPDS@CAR-M (ns: no significance, N = 3). (F) TEM images showing the morphology of mPDS, CAR-M, and mPDS@CAR-M (scale bar: 200 nm, 100 nm). (G) In vitro biocompatibility of mPDS@CAR-M, demonstrating no significant hemolysis compared to the positive control group with water (N = 3). (H) Drug release profiles of DOX and SOR from mPDS@CAR-M under NIR irradiation and non-irradiated conditions. (I) SDS-PAGE analysis of membrane proteins derived from K1 cells (lane 1), mPDS@M (lane 2), CAR-K1 cells (lane 3), and mPDS@CAR-M (lane 4). (J) Cellular uptake of mPDS@CAR-M observed by TEM (scale bar: 1 µm, 500 nm, 100 nm).

The preparation of mPDS@CAR-M involved three sequential steps:

(1) isolating plasma membrane vesicles (CAR-M) from CAR-K1 cells after removal of organelle membranes; (2) loading DOX and SOR into the mesoporous structure of mPDA to construct nanoparticles (mPDS); (3) mixing CAR-M with mPDS, followed by sequential co-extrusion of the mixture through 400 nm and 200 nm polycarbonate membranes to encapsulate mPDS within the membrane vesicles.

Dynamic light scattering (DLS) analysis revealed that mPDS@CAR-M exhibited an average diameter of approximately 190 nm, with the surface potential largely restored to the level of CAR-M (Figure 2E). The surface structure of mPDS was observed through SEM (Figure 1B), while TEM (Figure 2F) confirmed the morphological characteristics of mPDS, CAR-M, and mPDS@CAR-M. Additionally, the biological TEM images demonstrated the cellular uptake behavior and the distribution of mPDS@CAR-M within the cells (Figure 2J). Notably, the obtained mPDS@CAR-M maintained structural stability for at least two days (Supplementary Figure 1C and E), ensuring the technical feasibility of subsequent experiments.

Infrared and UV–visible absorption spectra of mPDS confirmed the successful loading of DOX and SOR into mPDA (Supplementary Figure 1D). Drug-loading efficiency of DOX was quantified by UV absorption analysis, while that of SOR was determined using high-performance liquid chromatography (HPLC). Based on standard calibration curves (Supplementary Figure 1F), the drug-loading efficiency and encapsulation efficiency of DOX in mPDS were calculated as 15.9% and 37.7%, respectively, whereas those of SOR were 16.2% and 38.8%. For single-drug formulations, the loading and encapsulation efficiencies of DOX in mPD were 18.2% and 44.4%, respectively, and those of SOR in mPS were 19.3% and 47.9% (Supplementary Table 2).

The biocompatibility of mPDS@CAR-M was evaluated at five concentrations (10, 30, 100, 300, and 1000 μg/ml). No significant hemolysis was observed in the mPDS@CAR-M groups compared with the positive control (Figure 2G). Furthermore, neither mPDA, mPDA@CAR-M, nor NIR irradiation alone showed any cytotoxic effects (Supplementary Figure 1I).

The in vivo photothermal effect forms the theoretical basis of photothermal therapy, in which elevated temperatures of 46–52 °C induce microvascular thrombosis, ischemia, and rapid cell death[12]. We first assessed the in vitro photothermal performance of mPDS@CAR-M. Upon 808 nm NIR laser irradiation, the temperature of the mPDS@CAR-M suspension reached 55 °C within 2 minutes and returned to room temperature approximately 2 minutes after laser withdrawal, demonstrating reproducible and stable photothermal behavior (Supplementary Figure 1G and H). The in vivo photothermal effect of mPDS@CAR-M was further confirmed (Figure 3G). In the absence of NIR irradiation, the vesicular coating effectively blocked mPDA, thereby restricting drug release from the nanoparticles[13], the cumulative release of DOX and SOR did not exceed 30% within 48 hours. In contrast, NIR irradiation induced vesicle rupture due to localized heating and increased the kinetic energy of drug molecules in solution[14]. The cumulative release of both drugs increased markedly within the first 8 hours and plateaued thereafter at approximately 80% (Figure 2H).

**Figure 3.**
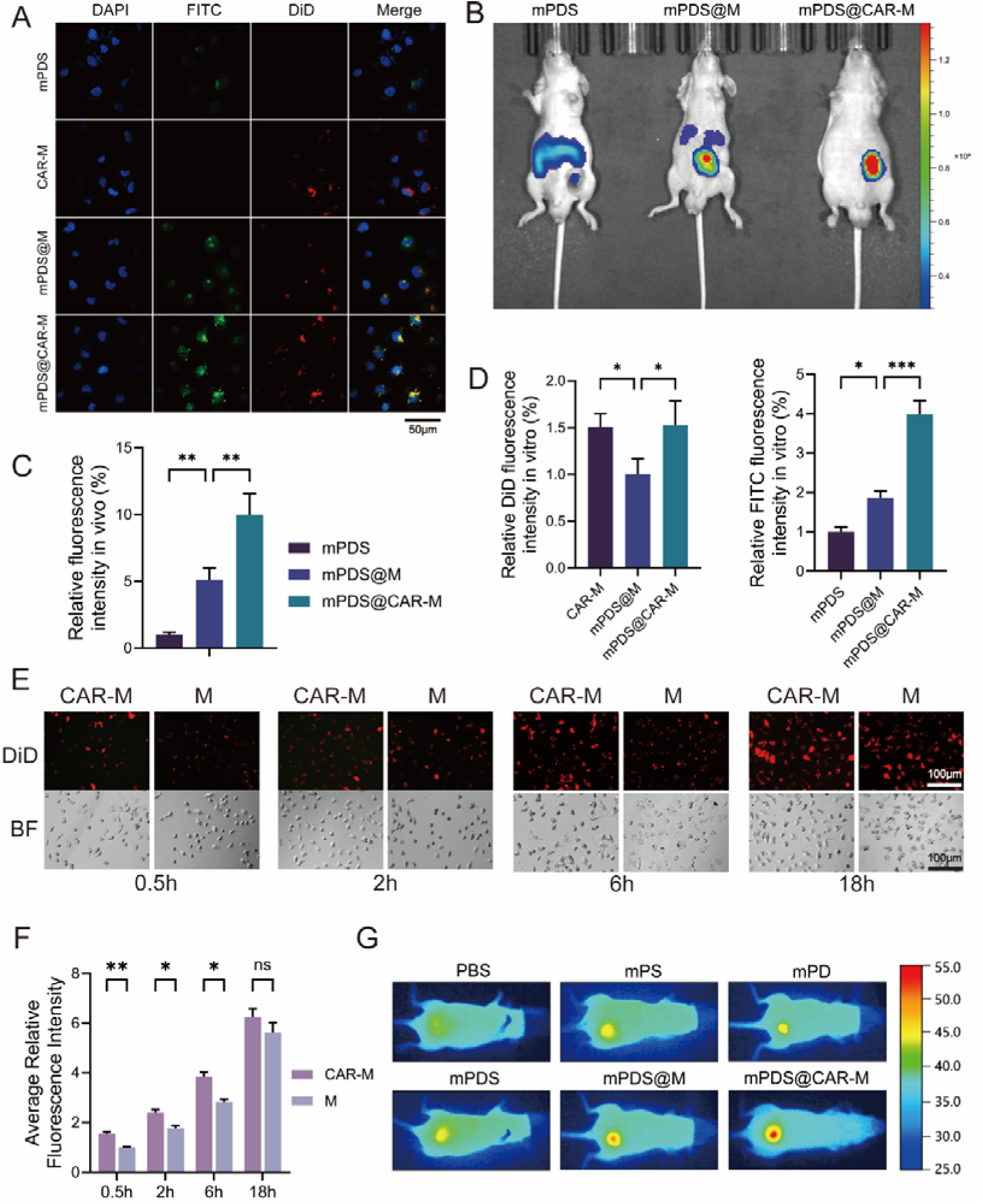
In vitro and in vivo tumor targeting of mPDS@CAR-M. (A) mPDS was labeled with FITC (green) and the membrane was labeled with DiD (red). The tumor cell-targeting ability of mPDS@CAR-M was investigated in vitro (scale bar: 50 μm). (B, C) Tumor accumulation of mPDS@CAR-M was evaluated using a xenograft tumor model in vivo (**p < 0.01, N = 3). (D) Statistical analysis of the fluorescence intensity of DiD and FITC in different groups in vitro (*p < 0.05, ***p < 0.001, N = 3). (E) Binding of CAR-M and M to K1-TSHR cells at different time points (scale bar: 100 μm) and (F) statistical analysis (ns: no significance, *p < 0.05, **p < 0.01, N = 3). (G) The photothermal effect of mPDS@CAR-M under 808 nm NIR irradiation in vivo.

Sodium Dodecyl Sulfate–Polyacrylamide Gel Electrophoresis (SDS-PAGE) revealed that mPDS@M and mPDS@CAR-M shared nearly identical membrane protein compositions with their corresponding vesicles (Figure 2I). Conserved adhesion molecules facilitated recognition and homotypic targeting of tumor cells through the formation of adhesive junctions[15,16], while CD47 functioned as a “self-marker” to protect against macrophage-mediated phagocytosis[17].

We evaluated the tumor-targeting ability of mPDS@CAR-M both in vitro and in vivo. mPDS was labeled with FITC (green), while the cell membrane was labeled with DiD (red). After co-incubating K1-TSHR cells with mPDS, CAR-M, mPDS@M, or mPDS@CAR-M, drug uptake was assessed by confocal microscopy. The mPDS@CAR-M group exhibited the strongest green fluorescence signal, with no significant difference in red fluorescence compared to the CAR-M group (Figure 3A and D), indicating that the in vitro tumor-targeting capability of mPDS@CAR-M was entirely derived from CAR-M. The tumor accumulation capacity of nanodrugs was further investigated in vivo using a xenograft tumor model and FITC-labeled mPDS. Nude mice were intravenously injected with mPDS, mPDS@M, or mPDS@CAR-M, and fluorescence intensity was measured 24 hours later using an in vivo imaging system. The fluorescence intensity at the tumor site in the mPDS@CAR-M group was significantly higher than that in the other groups (Figure 3B and C). Meanwhile, the accumulation in the liver, spleen, and kidneys was significantly lower than that in the other groups (Supplementary Figure 2A and B). This confirms that mPDS@CAR-M exhibits effective tumor-targeting ability both in vitro and in vivo.

We aimed to evaluate the relationship between the targeting ability of CAR-M and time. K1-TSHR cells were treated with DiD-labeled M or CAR-M. At 0.5 hours post-treatment, a higher signal of CAR-M was detected on the cell membrane of K1-TSHR cells compared to the M group (Figure 3E and F). This increased adhesion capability persisted up to 6 hours post-treatment. At 18 hours, no significant difference in fluorescence intensity was observed between the CM-DiD and CAR-M-DiD groups. The specific affinity of CAR-M was further evaluated in vitro by co-culturing with K1-TSHR and wild-type K1 cells (Supplementary Figure 2C and D). At different time points, K1-TSHR cells showed higher uptake efficiency and stronger fluorescence intensity compared to K1 cells.

We next investigated the antitumor effects of mPDS@CAR-M in thyroid cancer cells. Tumor cells were incubated with different concentrations of PBS, mPS, mPD, mPDS, mPDS@M, or mPDS@CAR-M for various durations. CCK-8 assays demonstrated that, compared with mPDS@M, mPDS@CAR-M further reduced cell viability after 48 h of treatment (Figure 4B), exhibiting both time- and concentration-dependent cytotoxicity. Based on these findings, 48 h and 10 μg were selected as the parameters for subsequent functional assays. Live/dead staining (Figure 4A and D), colony formation assays (Figure 4C and D), and Transwell assays (Figure 4E and D) collectively demonstrated the superiority of mPDS@CAR-M in terms of cytotoxicity, inhibition of colony formation, and suppression of cell migration, respectively, confirming its potent in vitro antitumor activity. These findings were further validated in IHH4-TSHR cells, yielding consistent results (Supplementary Figure 3A-E).

**Figure 4.**
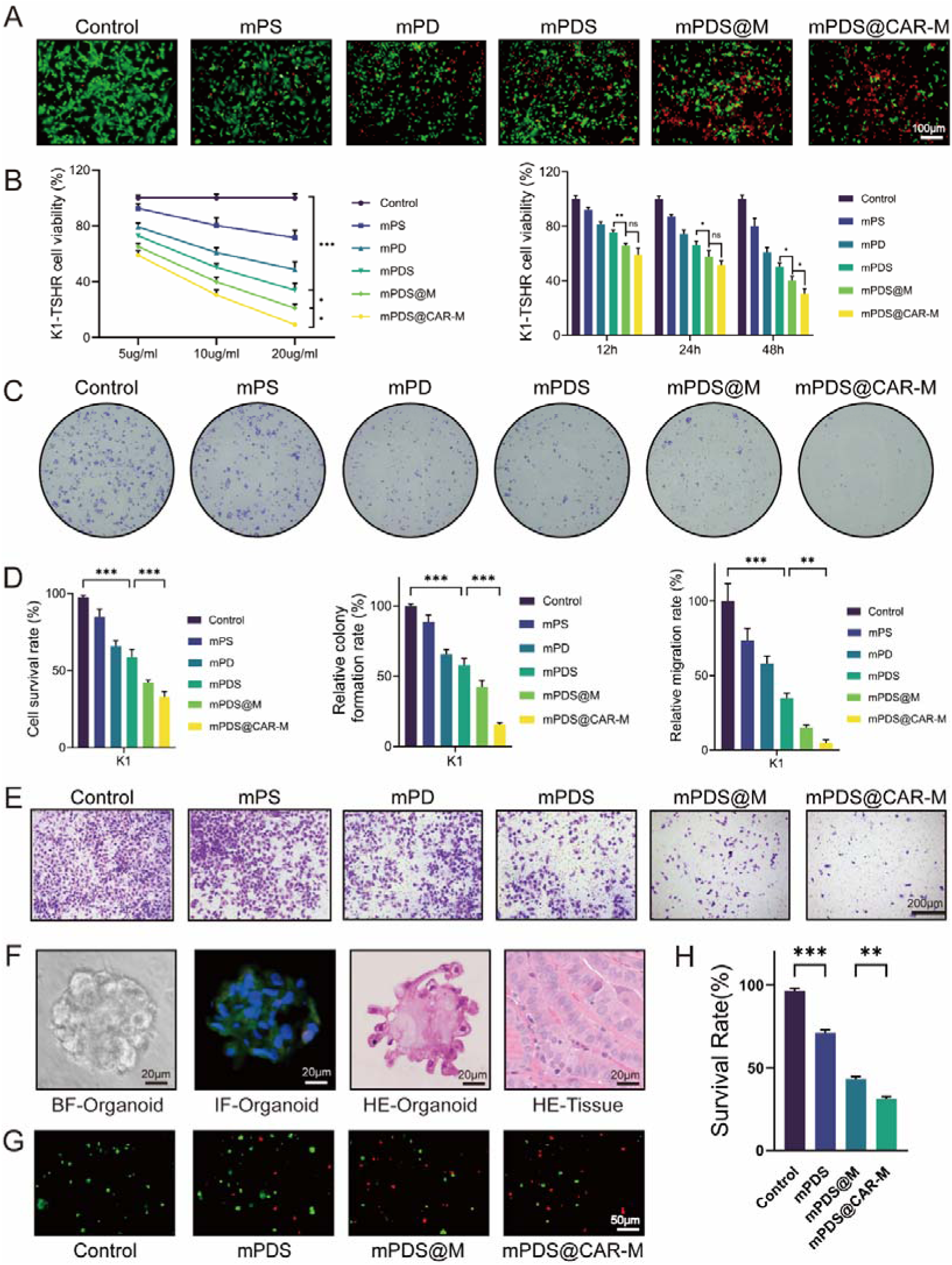
Antitumor efficacy of mPDS@CAR-M in vitro. (A) Live/dead staining assay demonstrating the effect of mPDS@CAR-M on K1-TSHR cells death (scale bar: 100 μm). (B) CCK-8 assay assessing the effect of mPDS@CAR-M on K1-TSHR cells viability (ns: no significance, *p < 0.05, **p < 0.01, ***p < 0.001, N = 3). (C) Colony formation assay showing the impact of mPDS@CAR-M on colony formation of K1-TSHR cells. (D) Statistical analysis of live/dead staining, colony formation, and transwell assays (**p < 0.01, ***p < 0.001, N = 3). (E) Transwell assay evaluating the effect of mPDS@CAR-M on K1-TSHR cells migration (scale bar: 200 μm). (F) Light microscopy images of thyroid cancer organoids, H&E staining, immunofluorescence, and corresponding H&E-stained tumor tissue sections (scale bar: 20 μm). (G) Antitumor effect of mPDS@CAR-M in thyroid cancer organoids and (H) quantitative analysis (**p < 0.01, ***p < 0.001, N = 3).

Subsequently, we assessed the therapeutic potential of nanomedicines in organoid models derived from thyroid cancer patients. Based on previous reports[18], thyroid cancer organoid systems were successfully established from tumors surgically resected from patients, and passaged once the average size reached 100 μm (Supplementary Figure 4A). H&E staining was used to compare the histological features of the organoids with those of the corresponding primary tumors. Immunofluorescence staining revealed that the expression of TSHR (green) was well preserved in the organoids, while the expression of Ki-67 (red) was relatively weak (Figure 4F). Consistent with the cell assays, treatment with nanoparticle drugs showed that the mPDS@CAR-M group exhibited the strongest anti-tumor effect (Figures 4G and H). Additionally, the morphological changes of the organoids before and after treatment were shown under optical microscopy (Supplementary Figure 4B).

We further investigated the potential mechanisms underlying the inhibitory effects of mPDS@CAR-M on tumor growth and metastasis. RNA sequencing (RNA-seq) was performed on the K1-TSHR cell line from the control group (PBS) and the mPDS@CAR-M group immediately after treatment. The alignment rate of each sample to the reference genome exceeded 95% (Supplementary Table 3). Differential gene expression (DGE) analysis was performed using the DESeq2 algorithm, with a screening threshold of P-adjust < 0.05 and a fold change ≥ 2. A total of 3,274 differentially expressed genes (DEGs) were identified between the control and mPDS@CAR-M groups, comprising 2,227 upregulated and 1,047 downregulated genes (Figure 5A). A binary heatmap visualized these transcriptomic differences, with upregulated genes in red and downregulated in blue (Figure 5B). Kyoto Encyclopedia of Genes and Genomes (KEGG) pathway analysis revealed that the RAP1 signaling pathway and the PI3K-Akt signaling pathway were among the most significantly enriched pathways (Figure 5C). The RAP1 signaling pathway activates the PI3K-Akt pathway, thereby promoting the generation of its downstream effects[19]. Gene Set Enrichment Analysis (GSEA) similarly highlighted the enrichment of pathways associated with the regulation of PI3K activity (Figure 5E). Genetic alterations in the PI3K-AKT signaling pathway constitute a critical component of the pathogenesis of thyroid cancer and are fundamental driving factors in the development of RAIR-DTC[20]. Furthermore, combined KEGG and LogFC enrichment analysis demonstrated significant enrichment of autophagy-related pathways (Figure 5D). mTOR, located downstream of the PI3K/AKT signaling pathway, is closely associated with the regulation of autophagy through its phosphorylation[21,22]. A circular heatmap illustrates the expression of key genes in the RAP1-PI3K-autophagy pathway (Figure 5F).

**Figure 5.**
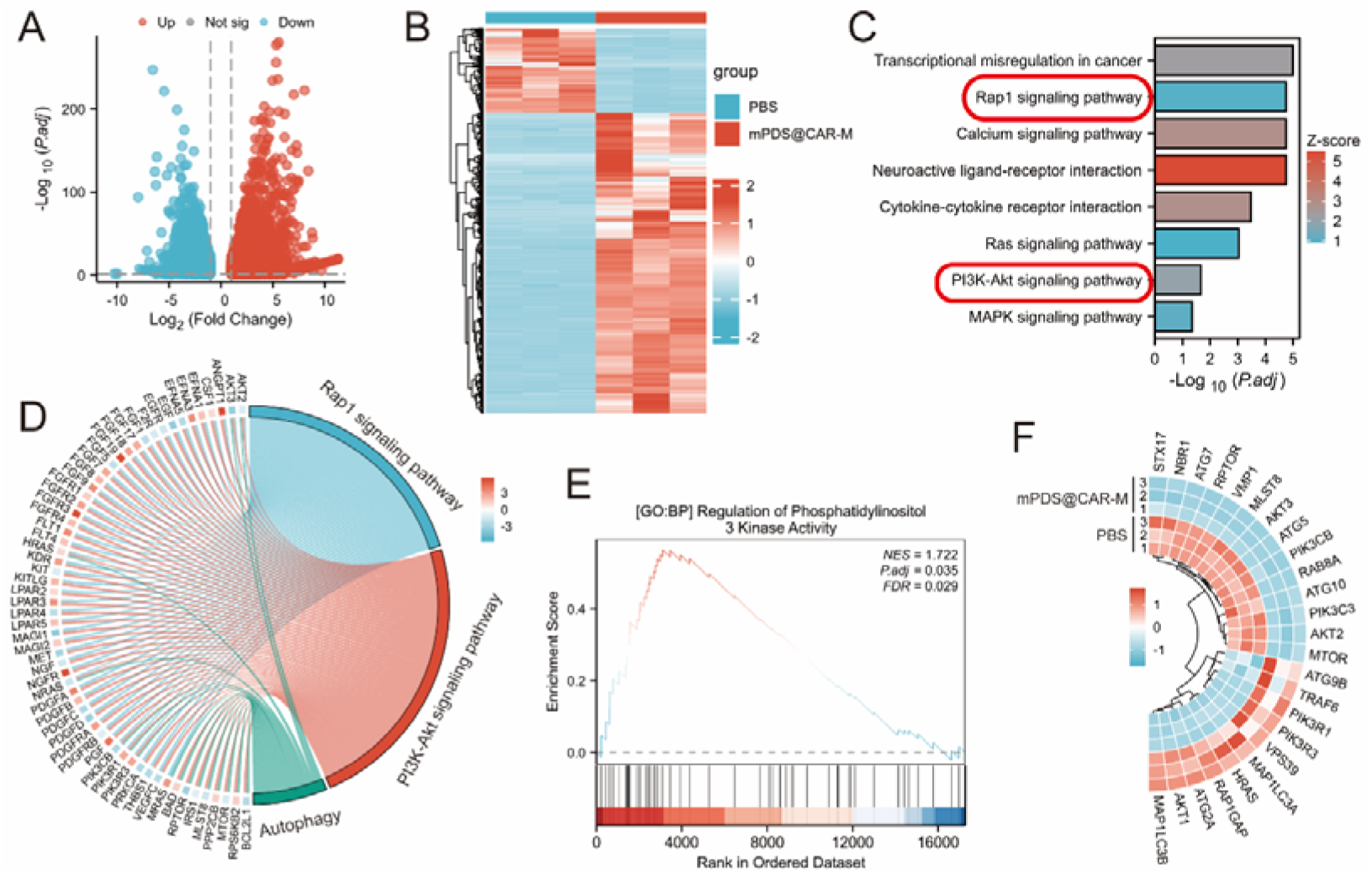
Transcriptome analysis of K1-TSHR cells treated with mPDS@CAR-M. (A, B) Volcano plot and heatmap of DEGs in K1-TSHR cells between the PBS group and the mPDS@CAR-M group. (C) GO and KEGG enrichment analysis of DEGs in K1-TSHR cells. (D) KEGG and LogFC-based combined enrichment analysis revealing significant enrichment of autophagy-related pathways. (E) GSEA identifying enrichment of PI3K activity-related pathways. (F) Heatmap of key genes in the RAP1-PI3K-autophagy pathway.

Intracellular glutathione (GSH) and reactive oxygen species (ROS) are essential regulators of redox homeostasis, as well as critical determinants of cell proliferation and death[23]. As a major intracellular antioxidant, GSH plays a central role in neutralizing ROS[24]. Studies have shown that polydopamine itself can lead to GSH depletion, possibly due to the antioxidant GSH promoting the depolymerization and degradation of mPDA[25]. Depletion of GSH results in excessive ROS accumulation, which induces oxidative stress, causes cellular damage, and disrupts key signaling pathways[26]. We observed that mPDS@CAR-M induced GSH depletion and ROS accumulation in a concentration-dependent manner (Figure 6A and B). GSH levels in tumor cells typically reach up to 10 mM, and it can synergistically accelerate the release of drug-loaded mPDA in the acidic microenvironment of tumor cells[27,28]. The released SOR, an xCT inhibitor, inhibits the activity of the SLC7A11 transporter (xCT), thereby suppressing the intracellular transport of GSH and exacerbating GSH depletion. We further validated the inhibitory effect of mPDS@CAR-M on xCT (Figure 6C). Meanwhile, the released DOX accumulates in the mitochondria, disrupting the respiratory chain and ultimately elevating ROS levels[29].

**Figure 6.**
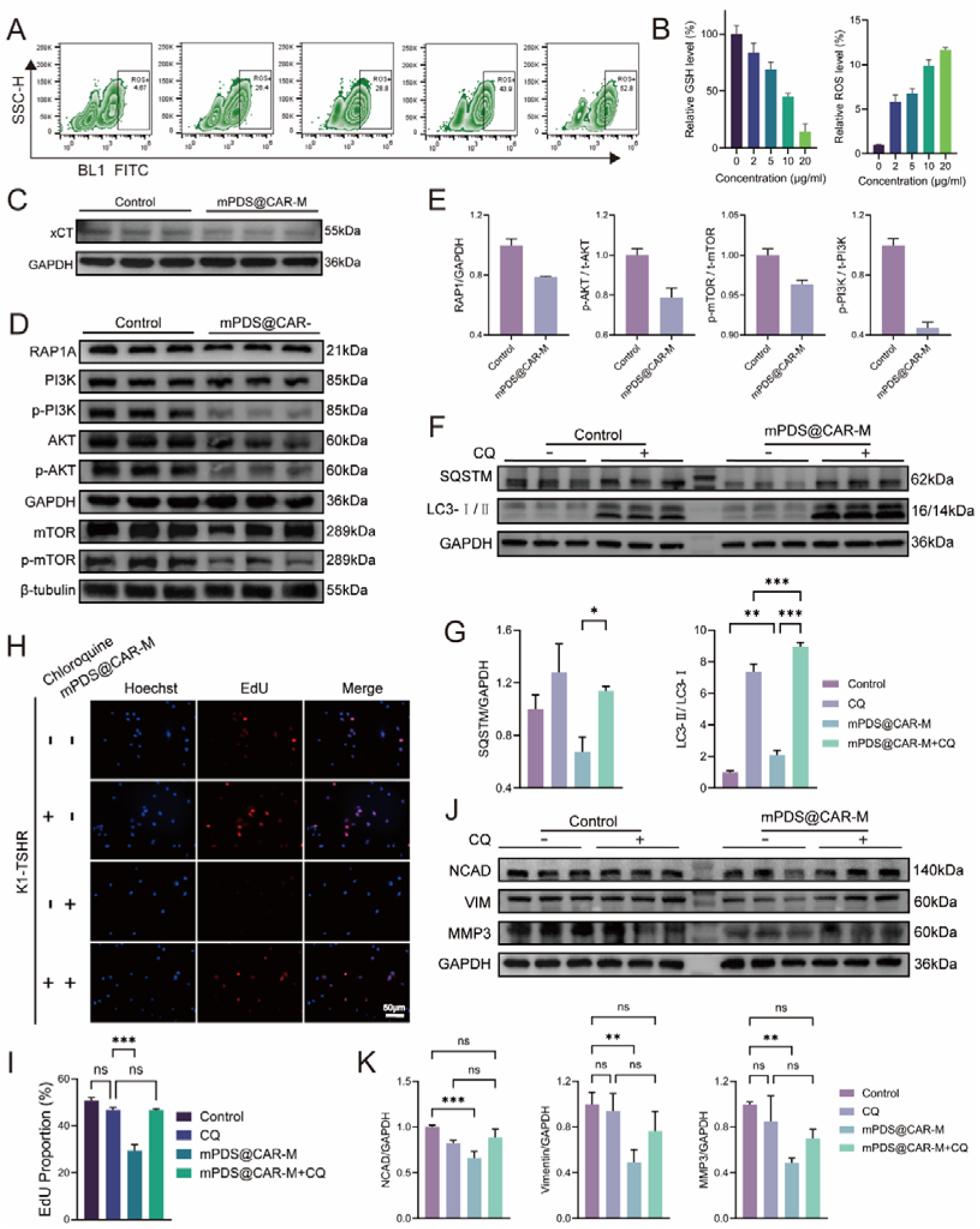
The potential mechanisms of mPDS@CAR-M therapy. (A) Flow cytometry analysis of intracellular ROS levels in cells treated with different concentrations of mPDS@CAR-M. (B) Quantitative analysis of the relationship between intracellular ROS and GSH levels and mPDS@CAR-M concentration (n = 3, mean ± SD). (C) Western blot analysis verifying the inhibitory effect of mPDS@CAR-M on xCT expression. (D) Western blot analysis of the expression levels of key proteins in the RAP1-PI3K-Akt-mTOR signaling pathway following mPDS@CAR-M treatment. (E) Quantitative analysis of the proteins in (D) (n = 3, mean ± SD). (F) Western blot analysis of autophagy-related protein expression after mPDS@CAR-M treatment. (G) Quantitative analysis of the proteins in (F) (n = 3, mean ± SD, *p < 0.05, **p < 0.01, ***p < 0.001). (H) EDU assay showing the reversal of mPDS@CAR-M’s antitumor effects by CQ on tumor cell proliferation (scale bar: 50 μm). (I) Quantitative analysis of the EDU assay results (n = 3, mean ± SD, ns: no significance, ***p < 0.001). (J) Western blot analysis of EMT-related protein expression after mPDS@CAR-M and CQ treatment. (K) Quantitative analysis of the proteins in (J) (n = 3, mean ± SD, **p < 0.01, ***p < 0.001).

We next examined key regulators of the RAP1–PI3K–Akt–mTOR signaling pathway by qPCR (Supplementary Figure 6A). Upregulation of RAP1GAP facilitated the conversion of RAP1-GTP to RAP1-GDP, thereby suppressed RAP1 signaling[30]. Reduced RAP1 activity in turn diminished its ability to recruit and activate PI3K, weakening the upstream signaling drive[31,32]. Downregulation of PIK3C3, PIK3CB, AKT1, AKT2, AKT3, MTOR, RPTOR, and MLST8 further suggested inhibition of the PI3K–AKT–mTOR pathway (Supplementary Figure 6A). Consistently, protein expression levels of p-PI3K, p-Akt, and p-mTOR were significantly lower in the mPDS@CAR-M group than in the control group (Figure 6D and E).

As a canonical regulatory axis of autophagy, inhibition of mTOR removes the brake on autophagy[33]. Western blotting (WB) confirmed the activation of autophagic flux (Figure 6F and G). Furthermore, rescue experiments demonstrated that the autophagy inhibitor chloroquine (CQ) completely reversed the antiproliferative effect of mPDS@CAR-M on tumor cells (Figure 6H and I), indicating that mPDS@CAR-M inhibit the proliferation of tumor cells through the activation of cell-damaging autophagy.

We also assessed epithelial–mesenchymal transition (EMT) related markers. The downregulation of FN1, VIM, CDH2, MMP2, MMP9, and SNAIL mRNA levels in the mPDS@CAR-M group compared to the control group (Supplementary Figure 6B), along with a significant decrease in the expression of EMT-related proteins NCAD, VIM, and MMP3 (Supplementary Figure 6C and D), suggests that mPDS@CAR-M effectively inhibits the EMT process, thereby suppressing tumor metastasis. Furthermore, when autophagy was inhibited by CQ, the expression of EMT-related proteins was restored to levels similar to the control group (Figure 6J and K). This suggests that mPDS@CAR-M inhibits EMT through the activation of detrimental autophagy.

To evaluate the in vivo antitumor efficacy of mPDS@CAR-M, a subcutaneous xenograft model was established in BALB/c nude mice using K1-TSHR cells. On day 10 after tumor implantation, mice received intravenous injections of the nanoparticle formulations via the tail vein, followed by NIR laser irradiation of the tumor site the next day (Figure 7A). The results showed that, compared to the control and mPDS groups, both the mPDS@M and mPDS@CAR-M groups exhibited significantly reduced tumor volumes (Figure 7B and C) and tumor weights (Supplementary Figure 5B), while no significant differences in body weight were observed among the groups (Figure 7D).

**Figure 7.**
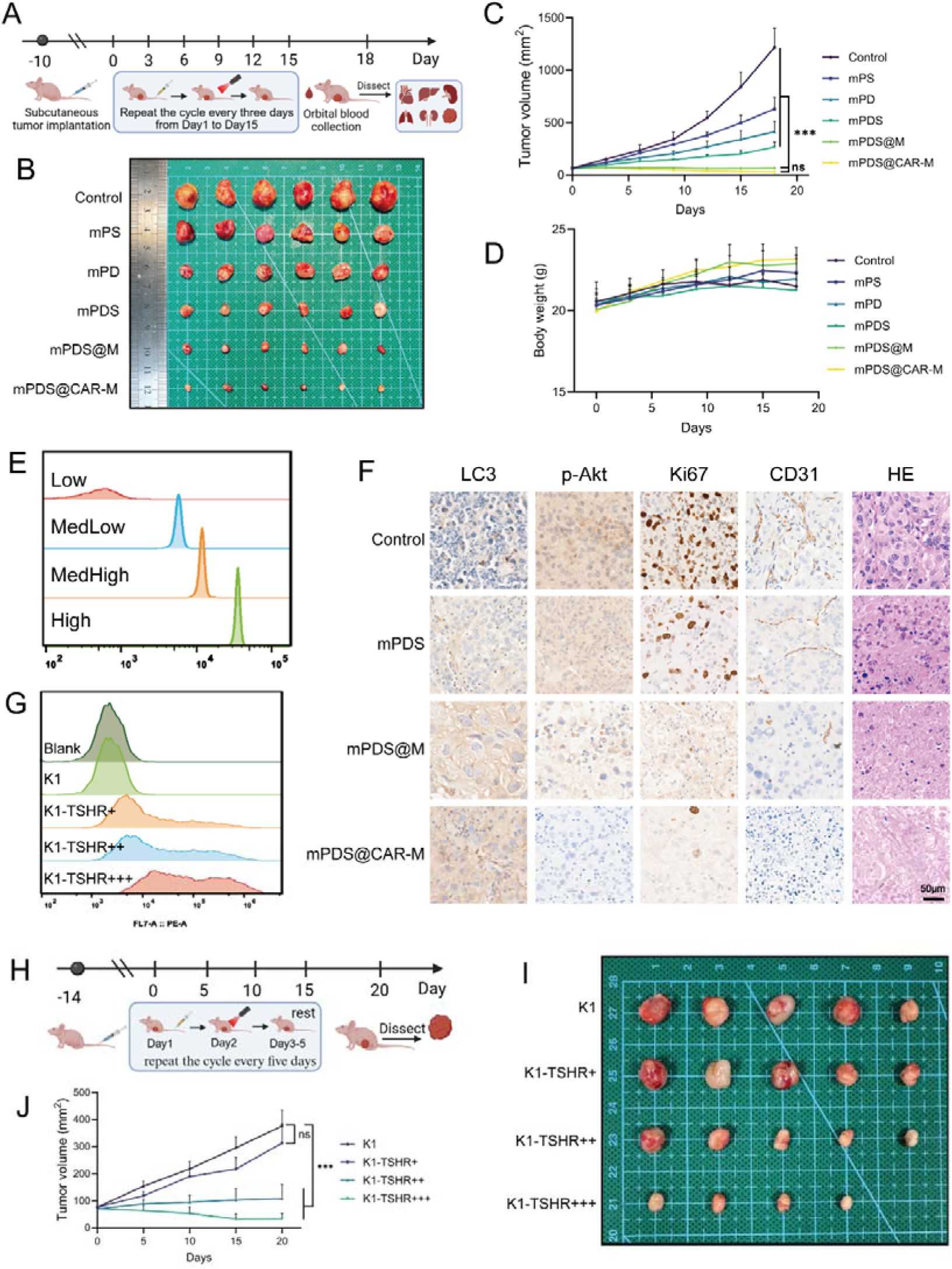
Targeted anti-tumor therapy of mPDS@CAR-M in vivo. (A) Schematic of the experimental timeline for tumor implantation, nanoparticle injection, laser irradiation, and tumor growth monitoring. (B, C) Nude mice were divided into groups: control, mPS, mPD, mPDS, mPDS@M, and mPDS@CAR-M. Tumor volumes were recorded over time (ns: no significance, ***p < 0.001, N = 6). (D) Monitoring of body weight in nude mice across above groups. (E) Standard curve for PE channel fluorescence intensity and the number of PE molecules bound to single cells. (F) H&E staining and IHC staining of Ki67, CD31, LC3, and p-Akt in control, mPDS, mPDS@M, and mPDS@CAR-M groups. (G) Quantitative analysis of TSHR expression levels on the surface of wild-type K1, K1-TSHR+, K1-TSHR++, and K1-TSHR+++ cells. (H) Experimental timeline for subcutaneous tumor formation in cells with varying TSHR expression levels, followed by mPDS@CAR-M injection, irradiation, and tumor growth monitoring. (I, J) Tumor growth curves and final volumes for subcutaneous tumors of K1, K1-TSHR+, K1-TSHR++, and K1-TSHR+++ cells (ns: no significance, ***p < 0.001, N = 5).

CK-MB and LDH1 are markers of myocardial injury, while AST and ALT serve as indicators of liver function, with ALT being more liver-specific and AST reflecting both hepatic and cardiac damage[34,35]. Following treatment, levels of CK-MB, LDH1, AST, and ALT in the mPDS@M and mPDS@CAR-M groups showed no significant differences compared with the control group, indicating favorable biosafety. In contrast, CK-MB and AST levels were significantly elevated in the mPDS group (Supplementary Figure 5A). Furthermore, hematoxylin and eosin (H&E) staining revealed no evident signs of cardiac damage, pulmonary toxicity, splenic inflammation, or hepatic and renal injury in the mPDS@M and mPDS@CAR-M groups (Supplementary Figure 5D).

H&E staining revealed disorganized tumor cell arrangement, cytoplasmic vacuolization, reduced nucleocytoplasmic contrast, and evidence of necrosis or fibrosis in the mPDS@M and mPDS@CAR-M groups. Immunohistochemical (IHC) analysis further demonstrated a marked reduction in Ki67-positive cells, indicating that mPDS@CAR-M exerted stronger antitumor proliferative in vivo. The expression of the neovascularization marker CD31 was significantly reduced in the mPDS@CAR-M group compared to the control group, which may be attributed to thermal ablation and the anti-angiogenic effect of sorafenib[36]. In the mPDS@CAR-M group, the number of p-Akt positive cells was significantly reduced, while the number of LC3 positive cells was markedly increased. This further confirms that mPDS@CAR-M kills tumor cells through the PI3K-Akt pathway and damage-induced autophagy (Figure 7F).

Although the antitumor effect of the mPDS@CAR-M group appeared stronger than that of the mPDS@M group (Figure 7B), no statistically significant differences in final tumor volume or weight were observed between the two groups (Figure 7C and Supplementary Figure 5B and C). We speculated that insufficient TSHR expression may have limited the in vivo targeting efficacy of the CAR structure. To investigate this, we generated cell lines with different levels of TSHR expression by transducing K1 cells with lentivirus at varying titers, designated as K1-TSHR+, K1-TSHR++, and K1-TSHR+++. Using calibration beads with four levels of PE conjugation, we established a standard curve that converted PE channel fluorescence intensity into the number of PE molecules bound per cell (Figure 7E). Quantitative analysis indicated that the relative surface expression levels of TSHR in K1-TSHR+, K1-TSHR++, and K1-TSHR+++ were approximately 1:2:6 (Figure 7G and Supplementary Table 4).

To evaluate the impact of TSHR expression on therapeutic efficacy, wild-type K1 cells and three engineered cell lines with varying TSHR levels were implanted in nude mice and treated with mPDS@CAR-M plus NIR irradiation (Figure 7H). Tumor size in the K1-TSHR++ group was significantly smaller than in the K1-TSHR+ group (Figure 7I and J), while no further reduction was observed in the K1-TSHR+++ group. These results suggest that a threshold level of TSHR expression is required for effective CAR-mediated targeting, and that excessively high expression does not confer additional therapeutic benefit. Given the heterogeneity of TSHR expression in patients, it is important to assess the TSHR expression levels[10].

## Conclusion

This study developed a genetically engineered cell membrane-coated nanodrug (mPDS@CAR-M), providing an innovative therapeutic strategy for RAIR-DTC. mPDS@CAR-M combines photothermal therapy, chemotherapy, and targeted therapy, effectively inhibiting tumor cell proliferation by enhancing oxidative stress, suppressing the PI3K-AKT-mTOR signaling pathway, and inducing cytotoxic autophagy. Additionally, mPDS@CAR-M activates harmful autophagy to inhibit EMT in tumor cells, thereby reducing their migratory capacity. In vivo experiments demonstrated that mPDS@CAR-M significantly reduced tumor volume, with its therapeutic efficacy being target number-dependent. In summary, mPDS@CAR-M exhibits considerable potential in the treatment of RAIR-DTC, and its clinical application warrants further exploration.

## Data Availability

The data supporting the conclusions of this study can be found within the article and its Supplementary Information files, and are also available from the corresponding author upon reasonable request.

## Credit author statement

Shaojie Xu, Xingyin Li and Youyun Peng contributed equally to this work.

Shaojie Xu, Xingyin Li and Youyun Peng: Writing – original draft, Methodology, Investigation, Formal analysis. Xue Yang: Supervision, Formal analysis, Data curation. Yuhang Su: Methodology, Formal analysis, Data curation. Sining Wang: Supervision, Data curation. Xingrui Li: Writing – review & editing, Supervision, Funding acquisition. Yaying Du: Writing – review & editing, Supervision, Funding acquisition.

