## Supplementary figures and images for "Genetically Engineered Cell Membrane-Coated Nanodrug for Targeted Treatment of Thyroid Cancer"

### Supplementary Figure 1

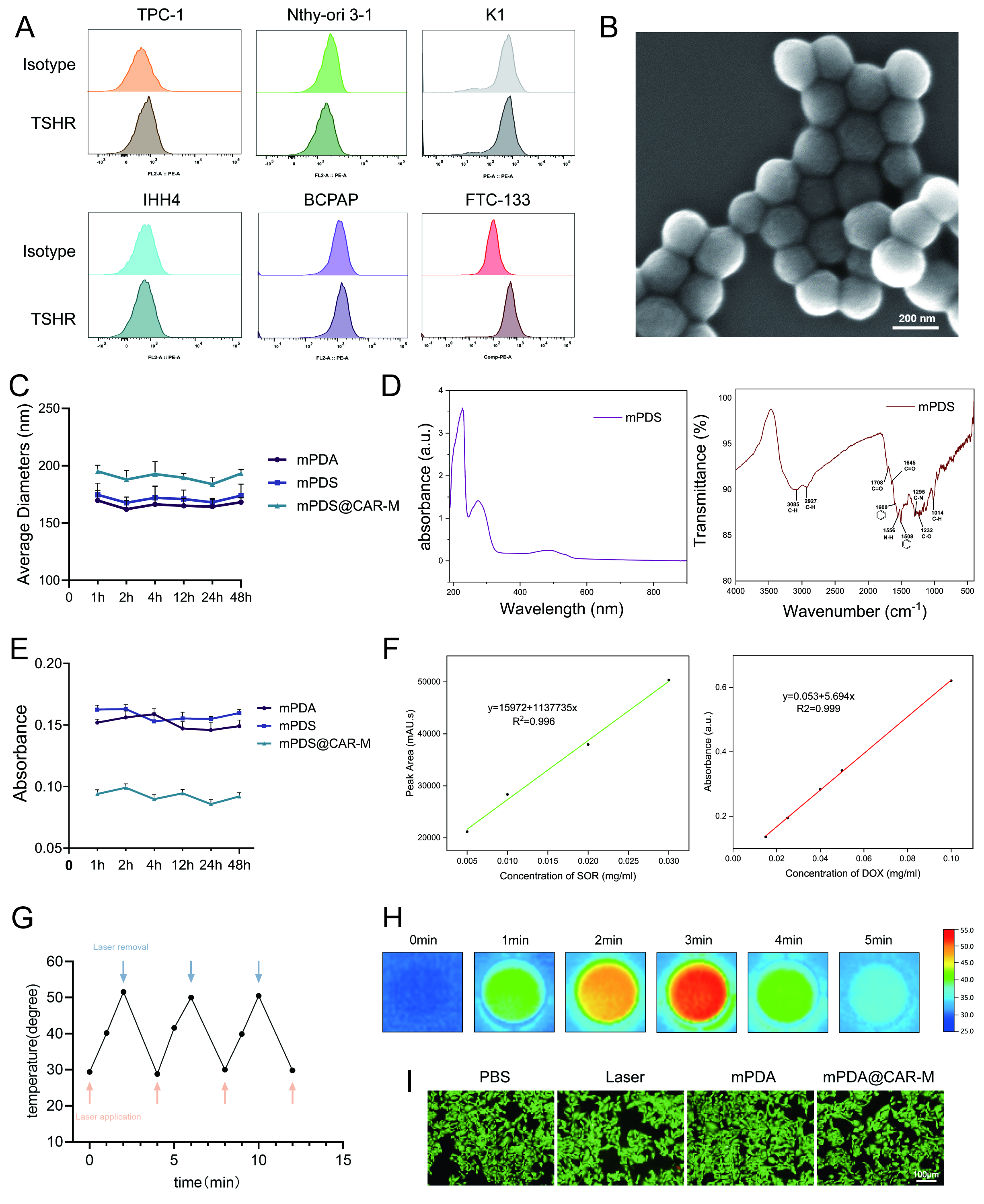

### Supplementary Figure 2

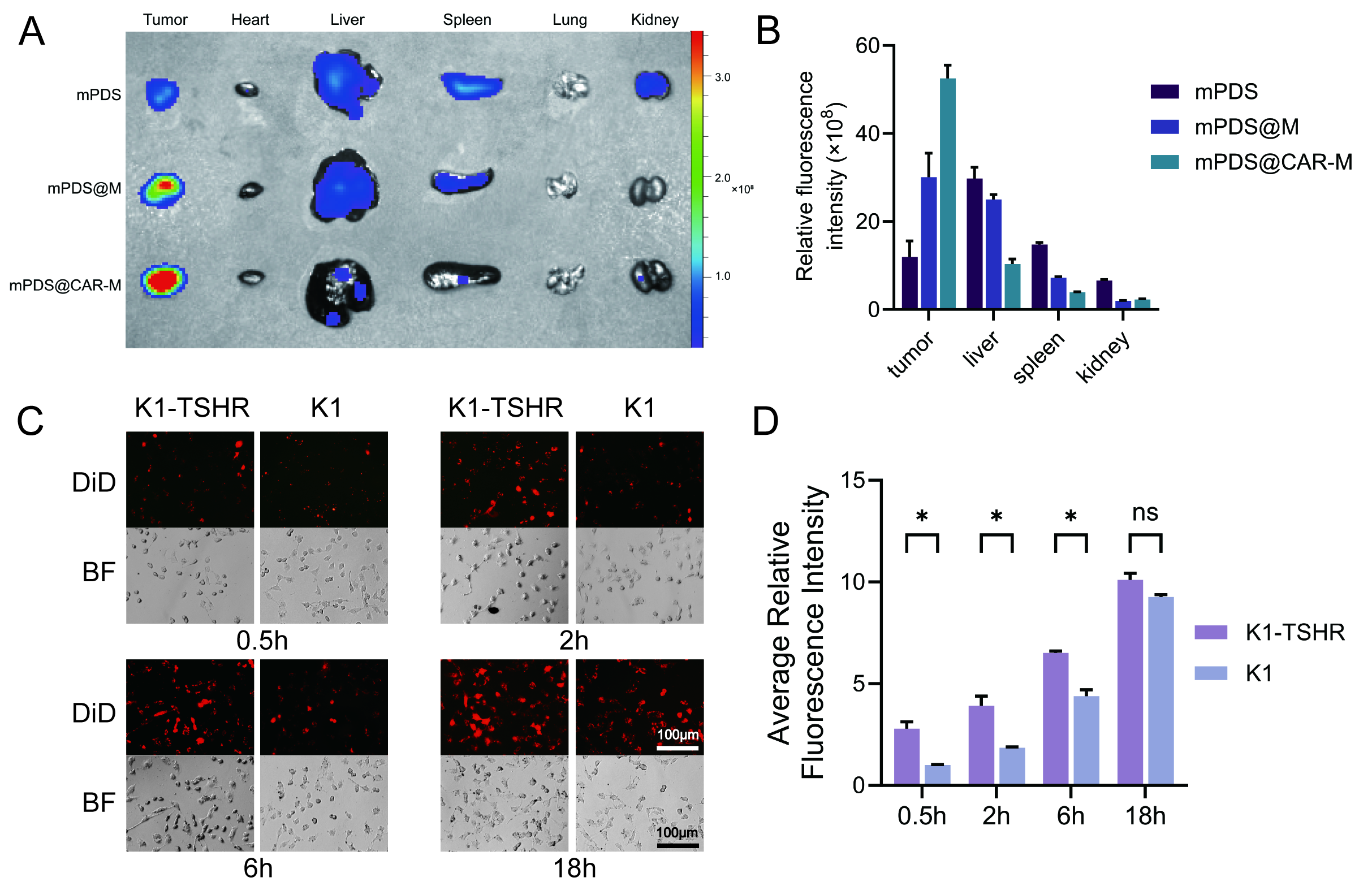

### Supplementary Figure 3

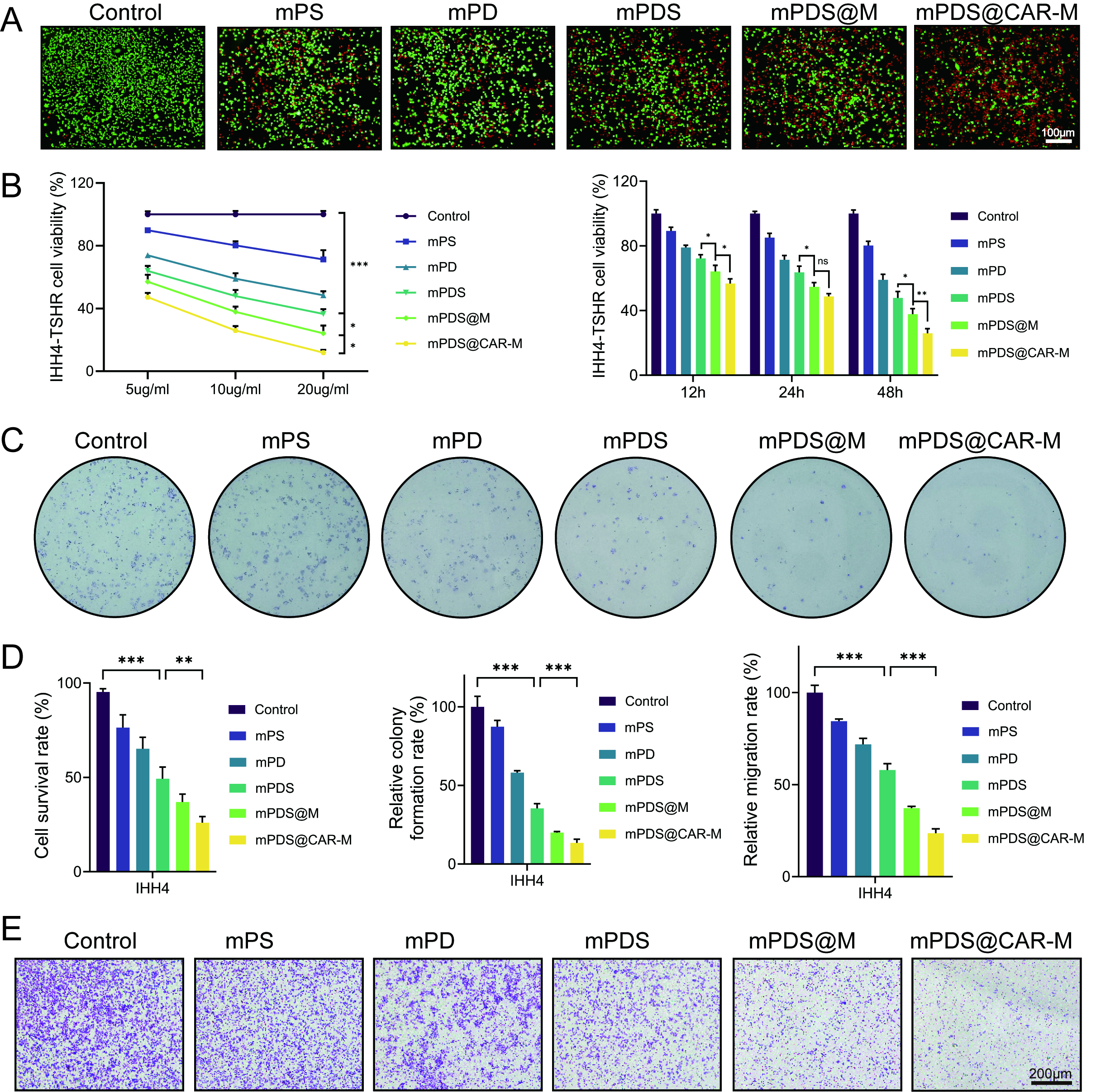

### Supplementary Figure 4

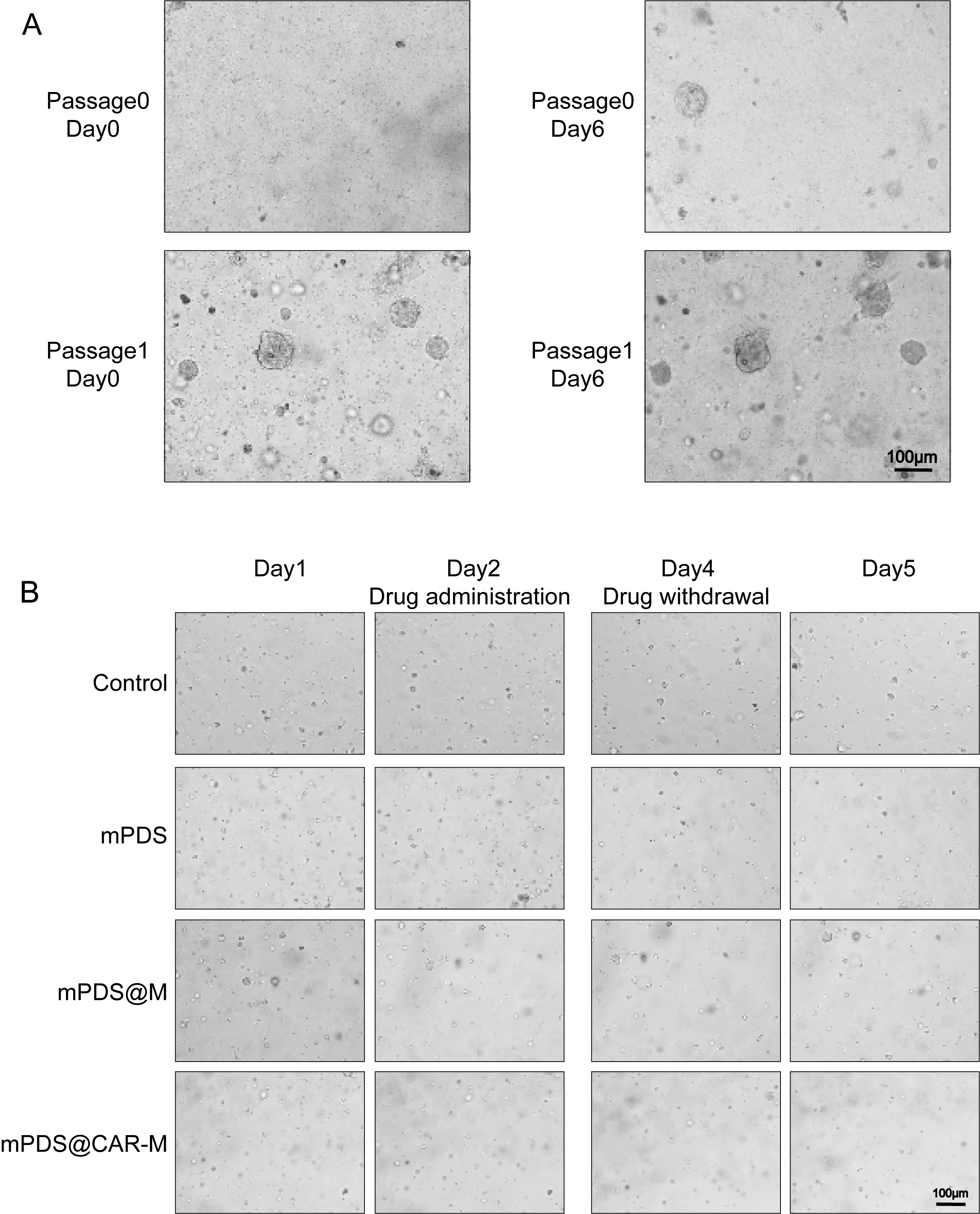

### Supplementary Figure 5

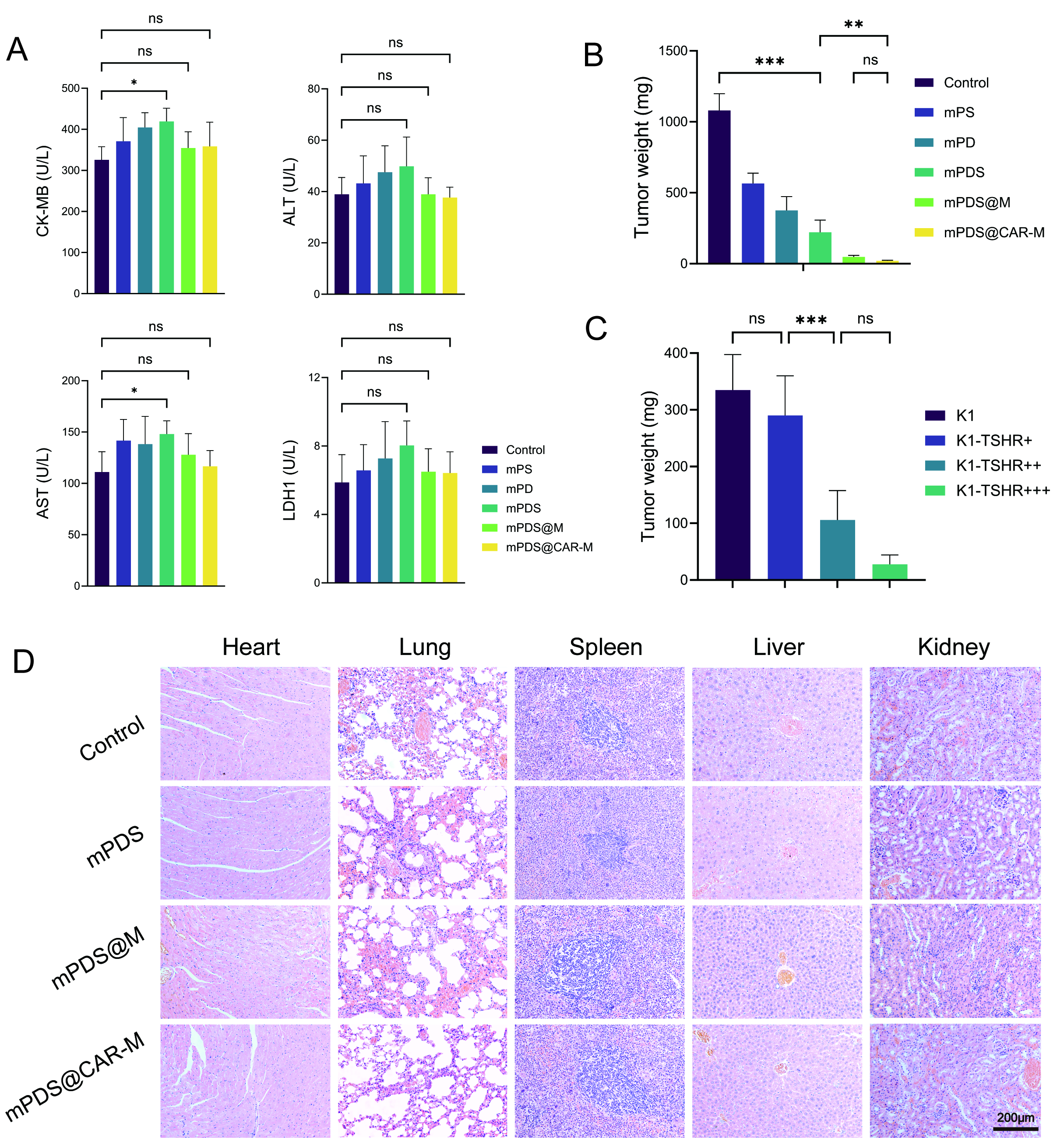

### Supplementary Figure 6

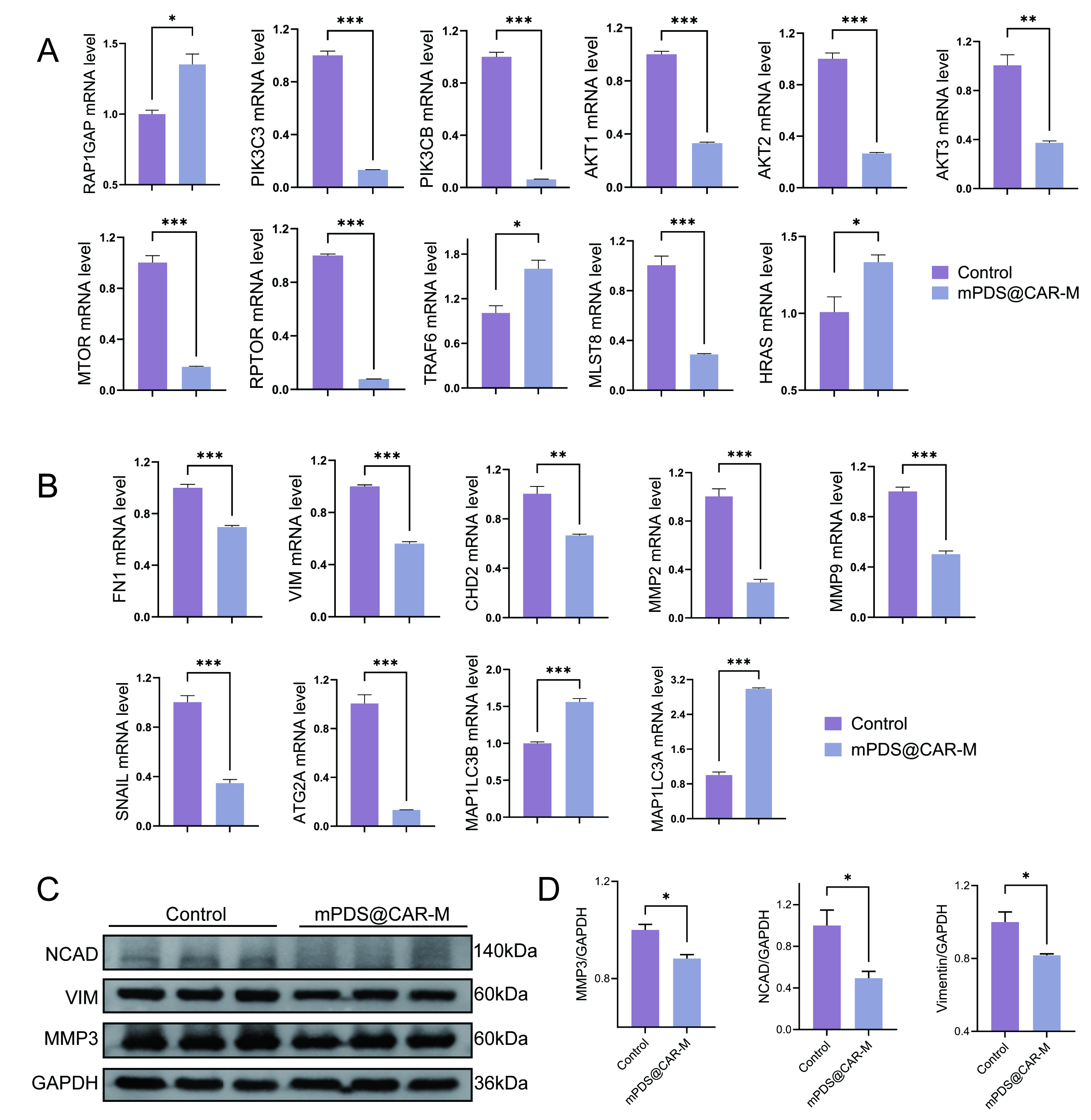
